# The accuracy and bias of single-step genomic prediction for populations under selection

**DOI:** 10.1101/090274

**Authors:** Wan-Ling Hsu, Dorian J. Garrick, Rohan L. Fernando

## Abstract

In single-step analyses, missing genotypes are explicitly or implicitly imputed, and this requires centering the observed genotypes, ideally using the mean of the unselected founders. If genotypes are only available on selected individuals, centering on the unselected founder mean is impossible. Here, computer simulation is used to study an alternative analysis that does not require centering genotypes but fits the mean *µ*_*g*_ of unselected individuals as a fixed effect. To improve numerical properties of the analysis, centering the entire matrix of observed and imputed genotypes, using their sample means can be done in addition to fitting *µ*_*g*_. Starting with observed diplotypes from 721 cattle, a 5 generation population was simulated with sire selection to produce 40,000 individuals with phenotypes of which the 1,000 sires had genotypes. The next generation of 8,000 genotyped individuals was used for validation. Evaluations were undertaken: with (J) or without (N) *µ*_*g*_ when marker covariates were not centered; and with (JC) or without (C) *µ*_*g*_ when all marker covariates were centered. A pedigree based evaluation was less accurate than genomic analyses. Centering did not influence accuracy of genomic prediction, but fitting *µ*_*g*_ did. Accuracies were improved when the panel comprised only QTL, models JC and J had accuracies of 99.2%; and models C and N had accuracies of 85.6%. When only markers were in the panel, the 4 models had accuracies of 63.9%. In panels that included causal variants, fitting *µ*_*g*_ in the model improved accuracy, but had little impact when the panel contained only markers.

---

In pedigree based analyses, the expected value of breeding values is zero. In order to achieve similar properties in whole-genome analyses, marker genotype covariates are often transformed. When all individuals are genotyped, it has been shown that inference on genotype effects does not depend on how the covariates are transformed (Strandén and Christensen 2010). However, when data includes genotyped and non-genotyped individuals, inference on marker effects from single-step analyses may depend on how the covariates are transformed. In single-step analyses using marker effects models, the breeding values of non-genotyped individuals are partitioned into components representing the prediction of non-genotyped individuals conditional on their genotyped relatives and an independent deviation (Fernando *et al*. 2014). The prediction of non-genotyped individuals conditional on their genotyped relatives is done based on best linear prediction, which requires the first moments to be known without error. This is straightforward if the mean of the genomic breeding value is zero in the absence of selection. Centering the observed genotype covariates using what their means would be in the absence of selection would result in genomic breeding values with null means. However, such genotype covariate means are typically unavailable. Fernando et al. (Fernando *et al*. 2014) proposed a solution for the marker effects model that involves fitting an additional fixed covariate that estimates the mean *µ*_*g*_ of the linear component of the genotypic value, which is denoted by *a_i_* in (1) below, in a population where selection is absent. Using that approach, even when there is selection, the selection process can be ignored (Goffinet 1983; Gianola and Fernando 1986; Im *et al*. 1989; Sorensen *et al*. 2001). In Markov chain Monte Carlo analyses, centering results in better mixing (Strandén and Christensen 2010), reducing the number of iterations required to obtain converged genomic predictions. In practice, centering the entire matrix of genotype covariates, including the observed and imputed genotypes, using their sample means can be done in addition to the Fernando et al. (Fernando *et al*. 2014) approach of fitting *µ*_*g*_. This type of centering of the entire genotype matrix does not affect inference on marker effects.

The same issue with centering of the observed genotype covariates that we described above for the marker effects model is also implicit for the single-step breeding value model (single step GBLUP), and a similar solution was proposed by Vitezica et al. (Vitezica *et al*. 2011). In their proposed solution, the observed genotype covariates are centered using their means, and in addition the genomic covariance matrix is corrected for the change in the mean breeding value of the genotyped individuals (Vitezica *et al*. 2011). It was shown in that paper that this is equivalent to fitting the change in breeding value due to selection as a random effect.

Here we use simulated data to compare the accuracy and bias in genomic prediction applied to populations under selection with and without centering the entire matrix of genotype covariates, and with and without fitting *µ*_*g*_ as a fixed effect. Further, we will show that when the observed genotype covariates are centered using means calculated from selected individuals rather than means from all individuals, the meaning of *µ*_*g*_ changes from the mean of unselected individuals to become the mean breeding value in selected individuals as claimed by Vitezica et al. (Vitezica *et al*. 2011).

## Materials and Methods

### Theory

To simplify the presentation of the genetic model, without loss of generality, we will assume that the unconditional expectation of the phenotypic value for all individuals is the same. Let 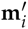 denote the row vector of genotypes for individual *i*. Then, under additive gene action, the genotypic value, *g_i_*, which is the expected phenotypic value of an individual with genotypes 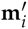 can be written as

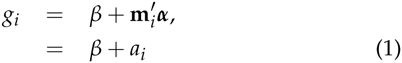

where *β* is the value of *g_i_* when 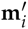 = **0**′ and ***α*** is the vector of substitution effects. Recognize the scalar *β* and the vector ***α*** are constants, but *g_i_* will be a random variable because of randomness in *a_i_* = 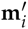***α***, due to the randomness in the genotypes for a randomly sampled individual. Note that the expected value of the linear component *a_i_* of the genotypic value in (1) is E(*a_i_*) = E(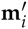) ***α*** = **k**′ ***α*** = *µ*_*g*_, where **k**′ = E(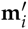), which may not be equal to zero. Thus, it is customary to write the model for the genotypic value, as can be derived from (1), as follows:

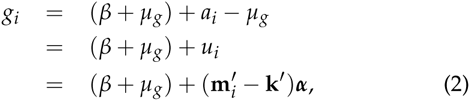

where (*β* + *µ*_*g*_) is a constant, representing the E(*g_i_*), and *u_i_* = (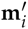 −**k**′) ***α*** is a random variable that has null expectation, which is the breeding value predicted in a pedigree-based BLUP evaluation. When genotypes are observed and used in a genomic analysis, they may be transformed or coded by subtracting their expectations, **k**′, from the observed values, 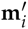. In both (1) and (2), *α_j_* is the same substitution effect for locus *j*. The intercepts in these models, however, are different. In (1), the intercept is *β*, and it is the value of *g_i_* when 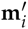 = **0**′. In (2), on the other hand, the intercept is (*β* + *µ*_*g*_), and it is the value of *g_i_* when 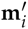 = **k**′.

More generally, **k**′ is not known, so genotypes are coded by subtracting a different vector **v**′ from the observed genotypes as 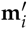 −**v**′. Still, *α_j_* is the substitution effect for locus *j*, but the intercept will change to become (*β* + **v**′***α***), which is the value of *g_i_* when 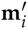 = **v**′. Thus, as more rigorously shown in (Strandén and Christensen 2010), inference about ***α*** does not depend on how the genotypes are coded. A simpler but rigorous proof is given in the appendix of this paper.

In single-step analyses, where some individuals are not genotyped, the missing genotypes are imputed either implicitly (Legarra *et al*. 2009) or explicitly (Fernando *et al*. 2014) using best linear prediction. Let **M**_*g*_ denote the matrix of genotypes for individuals that were genotyped. Then, the genotypes of the individuals with missing genotypes are imputed as

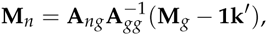

where **A***_ng_* is the matrix of pedigree based additive relationships between the non-genotyped and genotyped individual and **A***_gg_* is the matrix of additive relationships among genotyped individuals. Now, the model for the genotypic values, when genotypes are coded as in (2), becomes

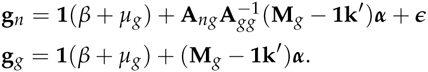

where ***∊*** is that part of **g***_n_* that cannot be imputed from knowledge of the breeding values of genotyped relatives. In practice, the true value of **k**′ is not known, and data for its estimation may not be available. Rearranging these equations in terms of the uncentered **M**_*g*_ rather than the centered matrix of geotype covariates (**M**_*g*_ − **1k**′), results in

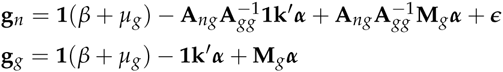

and substituting *µ*_*g*_ = **k**′***α***, as previously defined, results in

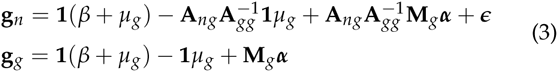

which suggests that *µ*_*g*_ = **k**′ ***α*** could be treated as an unknown constant and estimated as a fixed effect from the data (Fernando *et al*. 2014). The covariate vector for *µ*_*g*_ is denoted by **J***_n_* = 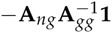 for non-genotyped individuals and by **J**_*g*_ = − **1** for genotyped individuals. So, (3) becomes

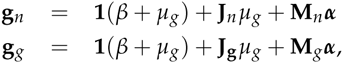

which can be combined as

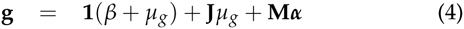

where 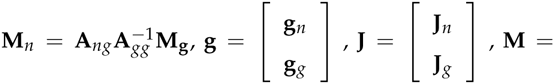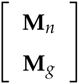.

When the vector ***α*** represents the substitution effects of a large number of loci containing positive and negative effects, *µ*_*g*_ = **k**′ ***α*** will tend to have a value close to zero. Accordingly, we have simulated some scenarios with positive *µ*_*α*_ = E(*α_i_*) so that the entire ***α*** vector is positive to exacerbate the impact of *µ*_*g*_ = **k**′ ***α***. Nevertheless, when marker rather than causal alleles are fitted in the model, the sign of the substitution effects depends on the phase relationship between marker and causal allele, which may be equally likely to be positive or negative.

Even if *µ*_*α*_ = *E(α_i_*) = 0, in a population undergoing selection, it is expected that E(*a_i_*) = E(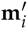) ***α*** ≠ 0 in non-founders. Suppose **v**′ is the mean of the observed genotype covariates in such a population undergoing selection and these means are used to center the matrix **M**_*g*_ of observed genotypes. Then, the model for the genotypic values can be written in terms of the matrix 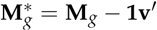 of centered covariates as

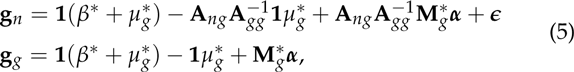

and using **J** for the covariate corresponding to *µ*_*g*_, (5) can be written as

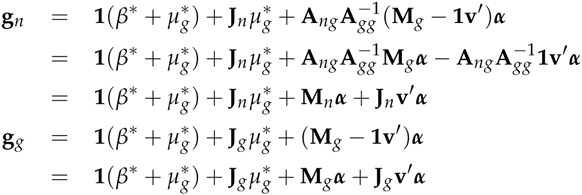

which can be combined as

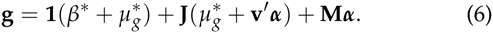

Note that the regression coefficients for **J**, *µ*_*g*_ in (4) and 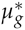 + **v**′ ***α*** in (6) must be equal. This implies that 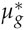 = *µ*_*g*_ − **v**′***α***. Similarly, the intercepts of these two models must be equal too, and this implies *β*\* = *β* + **v**′***α***.

### Simulations

Phenotypic and genotypic data were simulated based on haplotypes from ten regions of 721 US Hereford beef cattle that were genotyped with the Illumina 770K BovineHD BeadChip and reported in terms of the number of copies of the A allele at each locus. The selected regions came from choosing the 5,001st to 5,100th single nucleotide polymorphisms (SNPs) from chromosomes 1 to 10 (BTA1-BTA10), after eliminating SNP with MAF < 0.01. These remaining 1,000 SNP represent ten 0.1M chromosomes. Average LD between adjacent SNPs was 0.511. Half of these SNP were randomly chosen to represent QTL. The QTL effects were sampled from a Normal distribution with mean *µ*_*±*_ = 0.2 and multiplied by the number of copies of the A allele to produce the true breeding value (TBV). The TBV were added to a Normally distributed residual term scaled by the sample variance of the TBV to simulate a trait with a heritability of 0.5. The first 10 SNP (i.e. the 5,001st to 5,010th) from each of the ten chromosomes were also used to simulate a smaller panel, with 5 QTL and 5 markers per chromosome. Average LD between adjacent SNPs was 0.459. TBV were simulated in the same way as for the 1,000 SNP scenario, then scaled to simulate traits with heritabilities (*h*^2^) of 0.1, 0.3 or 0.5. An additional scenario with *µ*_*α*_ = 0 was used to simulate TBV for a trait with heritability 0.5.

Half the observed diplotypes from US Hereford cattle were assigned to represent males and the remainder to represent females. Those 360 males and 361 females were sampled in pairs, with replacement, to produce 4,000 male and 4,000 female offspring representing generation G-4. There were no mutations. Four more non-overlapping generations of random mating were carried out with one male and one female offspring per dam mated to randomly chosen sires to produce the founder population (G0).

The G1 generation was produced by mass phenotypic selection of the top 200 G0 males, and this was repeated for 5 more generations. Each female was randomly mated twice to selected males to produce 1 offspring of each sex each generation. Across non-overlapping generation G0 to G5, a total of 48,000 individuals with phenotypes, genotypes and TBV were simulated for each scenario.

The training data included phenotypes from all individuals in G0 to G4 (n = 40,000), and genotypes from all 1,000 sires and all 8,000 G5 animals. Fixed loci, if any, were filtered from the panel before genomic prediction analyses. The genetic and residual variances used in genomic prediction were the sample variance of the TBV in G0 and the corresponding residual variance used to define the desired heritability in the founder population.

### Models

Five statistical models were compared for differences in accuracy and bias of prediction. These include models with or without *µ*_*g*_ and with or without centering of marker covariates, and a model that used pedigree relationships but not marker covariates.

#### 1. Mixed Linear Model

Accuracy of pedigree-based best linear unbiased prediction (PBLUP) was quantified using the correlation of TBV and estimated breeding values (EBV), where TBV was as simulated and EBV were obtained by fitting the mixed linear model (Henderson 1973, 1984):

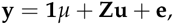

where **y** is a vector of phenotypic observations, **1** is a vector of 1s, *µ* = *β* + *µ*_*g*_ is a general mean, **u** is a vector of random direct additive genetic effects, **e** is a vector of random residual effects, and **Z** is a known incidence matrix relating observations to **u**. In this model, E(**u**) = 0, E(**e**) = 0, so that E(**y**) = **1***µ*. Further, 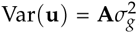, for **A** being the numerator relationship matrix, and 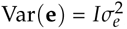, so that 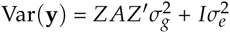.

#### 2. Single-Step Bayesian Regression Model

Genomic EBVs (GEBV) were obtained by Single-Step Bayesian regression (Fernando *et al*. 2014) with BayesC priors for marker effects with *π* = 0. The model was implemented in Julia (http://julialang.org) based on the SSBR package (http://QTL.rocks) to construct an MCMC chain of 50,000 samples. Individuals were separated into 2 groups designated with subscripts *g* or *n* according to whether or not simulated genotypes were assumed to be observed or missing. The single-step bayesian regression model including a covariate **J** for *µ*_*g*_ (Model J) was:

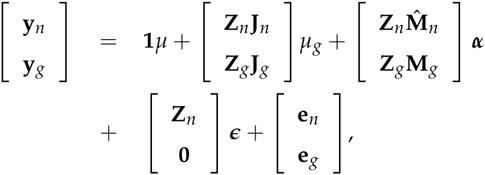

where **y***_n_* and **y**_*g*_ are vectors of phenotypes for nongenotyped and genotyped individuals, **1** is a vector of 1s, *µ* is a general mean, *µ*_*g*_ is the expected value of the linear component *a_i_* of the genotypic value if selection was absent, ***α*** is a vector of random substitution effects of markers, ***∊*** a vector of imputation residuals, **Z***_n_* and **Z**_*g*_ are incidence matrices relating the breeding values of non-genotyped and genotyped individuals to their phenotypes, **J**_*g*_, which is defined for genotyped individuals, is a vector of −1s, **J***_n_*, which is defined for non-genotyped individuals, is a vector computed as 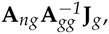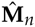 is the matrix of imputed marker covariates, **M**_*g*_ is the matrix of observed marker covariates, **e***_n_* and **e**_*g*_ are vectors of random residual effects for non-genotyped and genotyped individuals. This model can be represented as

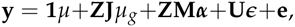

where 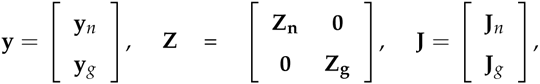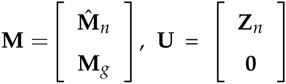, the **0** matrix in **U** is required because ***ϵ*** does not appear in the model for genotyped individuals, and **e** is a vector of random residual effects.

There were four variants of the single-step Bayesian analysis depending on whether or not the covariate **J** corresponding to the mean *µ*_*g*_ was in the model, and whether or not the columns in the marker covariate matrix **M** were centered using their observed means. The analyses with **J** or without **J** are are denoted as J or N when covariates were not centered, and as JC or C, when the entire matrix of imputed and observed genotype covariates were centered, respectively (Table 1).

**Table 1.**
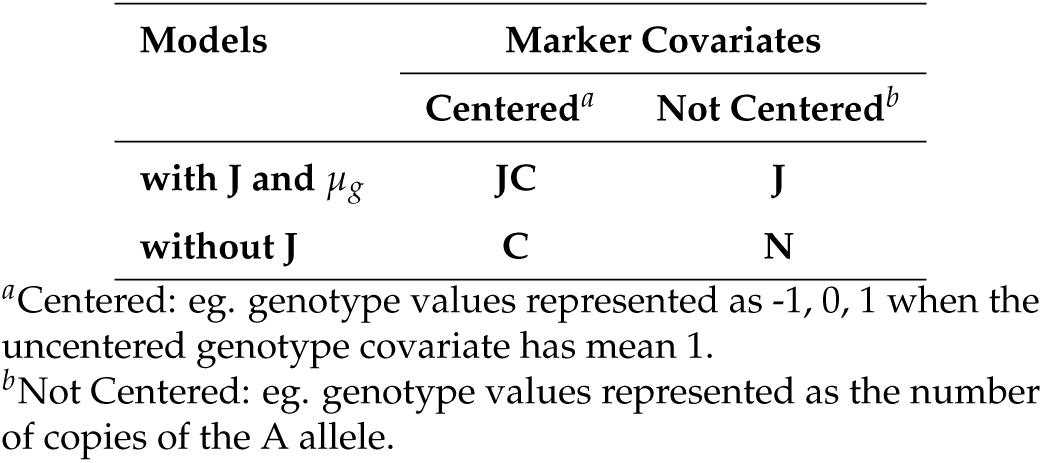
Four combinations of the single-step bayesian regression analyses

| Models | Marker Covariates |  |
| --- | --- | --- |
|  | Centered<sup>a</sup> | Not Centered<sup>b</sup> |
| with J and $\mu_g$ | JC | J |
| without J | C | N |

<sup>a</sup>Centered: eg. genotype values represented as -1, 0, 1 when the uncentered genotype covariate has mean 1.
<sup>b</sup>Not Centered: eg. genotype values represented as the number of copies of the A allele.

Accuracy of genomic prediction was quantified using the correlation of TBV and GEBV (*r*_*g*_,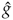), where GEBV were obtained from each of the 4 analyses described above. Bias of genomic prediction was quantified using the deviation from unity of the coefficient of regression of TBV on GEBV (*b*_*g*_,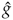). In models JC and J, GEBV are obtained using equation (24) in (Fernando *et al*. 2014):

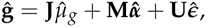

where 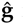 is the GEBV, 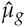 is the best linear unbiased estimate of the mean of breeding values, 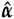 is the best linear unbiased predictor (BLUP) of the vector of random substitution effects of all markers, and 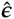 is the BLUP of the imputation residual.

In model J the matrix **M**_*g*_ contains the uncentered number of copies of the A allele at each locus, and the uncentered version is used to impute 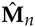. In model JC the entire matrix of imputed, 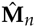, and observed, **M**_*g*_, genotype covariates is centered. In model C and N the GEBV are computed in a corresponding manner except that the covariate **J** and its coefficient *µ*_*g*_ are not included in the model.

The four analyses JC, J, C and N were all applied to 3 different genotype panels, comprising the causal QTL plus markers, just the causal QTL, or just the markers. All 12 combinations of 4 analyses and 3 genotype panels were applied to data simulated with 100 loci comprising 50 QTL whose effects were sampled from a Normal distribution with *µ*_*α*_ = 0.2 to construct phenotypes with *h*^2^ = 0.5. The four analyses JC, J, C and N were repeated using only genotype panels comprising QTL plus markers for four other scenarios: 100 loci, *h*^2^ = 0.1, *µ*_*α*_ = 0.2; 100 loci, *h*^2^ = 0.3, *µ*_*α*_ = 0.2; 100 loci, *h*^2^ = 0.5, *µ*_*α*_ = 0; and 1,000 loci, *h*^2^ = 0.5, *µ*_*α*_ = 0.2. Every scenario was repeated for 10 replicates with each replicate having been constructed starting from the sampling of G-5 which represented simulated offspring from the haplotypes of real animals. Every phenotypic dataset was also fitted using the PBLUP model. All reported correlations and regression coefficients are the means of 10 replicates. These are presented along with the standard errors of those means.

In single-step GBLUP (Legarra *et al*. 2009; Aguilar *et al*. 2010; Christensen and Lund 2010), the missing genotypes are not explicitly imputed and only the observed genotype covariates are centered using their means. So, in addition to the above, analyses with and without **J** (models JC* and C*), were applied to a marker panel with 100 loci, *h*^2^ = 0.5, and *µ*_*α*_ = 0, when the matrix 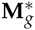 = **M**_g_ − **1v**′ of observed genotype covariates were centered using their means, **v**′ = **1**′**M**_*g*_. Recall that when the matrix 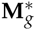 of observed genotype covariates is centered, the model for the genotypic values can be written in terms of the matrix **M**_*g*_ of uncentered covariates as shown by model (6). We will compare the estimates of 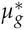 in this model with those of *µ*_*g*_ from (4) where the means of observed genotype covariates are not used for centering.

The genotypes representing G0 from each replicate and scenario are available at: https://figshare.com/s/d7798b811a9a6a4172fc. These genotypes and the methodology described previously are sufficient to reproduce the simulations used in this study.

## Results and Discussion

### Effect of fitting a genotyped mean and centering marker covariates

#### 1. Accuracy

The accuracies of genomic prediction as assessed by validation in G5 after training using G0-G4 for a trait with 50 QTL whose effects were sampled from a Normal distribution with *µ*_*α*_ = 0.2 and *h*^2^ = 0.5 are in Table 2. The accuracy of PBLUP in predicting breeding values for individuals without phenotypes, was 41.5%, accounting for less than 20% of genetic variance.

**Table 2.** Correlations (%, ±*SE*_*s*_) between TBV and (G)EBV^*a*^ for alternative analyses^*b*^

| Genotype Data<sup>c</sup> | Analyses |  |  |  |  |
| --- | --- | --- | --- | --- | --- |
|  | JC | J | C | N | PBLUP |
| 50 QTL + 50 Markers | 98.35 ± .00 | 98.41 ± .00 | 92.20 ± .00 | 92.18 ± .00 | - |
| 50 QTL Only | 99.19 ± .00 | 99.21 ± .00 | 85.61 ± .01 | 85.64 ± .01 | - |
| 50 Markers Only | 63.97 ± .02 | 63.96 ± .02 | 63.88 ± .02 | 63.88 ± .02 | - |
| No Genotypes | - | - | - | - | 41.53 ± .00 |

<sup>a</sup> Average correlation between true breeding value (TBV) and (genomic) estimated breeding values from 10 replications validated in Generation 5, comprising 8,000 individuals with genotypes but no phenotypes. The true QTL effects were sampled from a Normal distribution with mean $\mu_a = 0.2$ and scaled to simulate a trait with a heritability 0.5.
<sup>b</sup>J: includes a covariate for $\mu_g$ in the model, C: entire matrix of imputed and observed genotype covariates centered, JC: both J and C, N: neither J or C, and PBLUP: pedigree-based BLUP.
<sup>c</sup>The analyses were based on fitting covariates for only 50 QTL, only 50 markers, or both 50 QTL and 50 markers.

All analyses using genotypes resulted in more accurate predictions than using pedigree alone. Centering had no affect on the accuracy of the genomic analyses regardless of the nature of the marker panel. In this study, selection resulted in successive advance in mean TBV from G0 to G5 being 10.35, 10.77, 11.31, 11.82, 12.30 and 12.80. The mean genotypic value was not zero in G0 because QTL genotypes were not centered and the mean QTL effect was *µ*_*α*_ = 0.2. Since the QTL effects do not change by selection the advance in TBV reflects changes in the frequencies of the favorable alleles of the 50 QTL. So centering using the allele frequency means of the genotyped sires in G1-G4 and all individuals in G5 does not closely approximate the centering that would have occurred if the allele frequency means had been obtained from the unselected population. In contrast fitting *µ*_*g*_ in the model estimates the relevant mean from the data.

In panels that included causal variants (QTL), fitting *µ*_*g*_ in the model substantially improved the accuracy to being near perfect. This is not surprising given there were only 50 QTL, the heritability was 0.5 and there were 40,000 phenotyped ancestors, including 200 genotyped sires per generation in the training. However, in the panel that contained only markers with no causal variants, fitting *µ*_*g*_ in the model had little impact.

Using one replicate as an example, for the panel including both QTL and markers, the estimate of *µ* was about 10.51 for both analyses J and N. The estimate of *µ*_*g*_ was 7.66 for the analysis using J. For the genotyped individuals, the covariate values in **J**_*g*_ are all −1, so **1**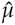 + **J**_*g*_ 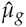 is a vector of values equal to 10.51 − 7.66 = 2.85. For non-genotyped individuals, the covariate values in **J***_n_* can vary widely but many are close to −1 while others are close to 0. This means that **1**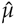 + **J***_n_* 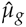 will contain values that range from 10.51 to 2.85, accounting for variation in accuracy of imputation. When *µ*_*g*_ is not included in the model, these effects are ignored which can reduce the accuracy of predicting nongenotyped individuals. Failing to account for these effects will propagate errors in 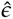 and 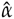, the latter impacting the accuracy of predicting genotyped individuals. Collectively, these errors reduced accuracy from 98% to 92% for the panel including QTL and markers and from 99% to 85% for the panel including only QTL. However, when the panel comprised only markers, the estimates 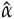 will include both positive and negative values because the phase of markers and QTL are equally likely to take either sign, in which case 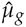 will be close to zero as confirmed in the above mentioned replicate where the estimate was 0.41.

Here, the QTL model was used with an intercept of *β* = 0.0 to simulate the data. When only QTL are on the panel, the true value of *β* is zero. Thus, in analysis J because *µ* = (*β* + *µ*_*g*_), both 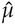 and 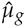 are estimates of *µ*_*g*_, and could be pooled which for the replicate above would be (10.51 + 9.04)/2 = 9.78. In that replicate, the actual mean of *a_i_* in G0 was 10.8, which was estimated in the analysis to be 9.78. On the other hand, the mean of the breeding value *u_i_* in the 9,000 genotyped individuals was 2.2, which is clearly not near the pooled estimate of 9.78. These genotyped individuals included 1,000 selected sires of which 200 were genotyped in each generation from G0 to G4, and 8,000 offspring from G5. The mean values of *u_i_* for the selected sires were 0.97, 1.41, 1.80, 2.24, and 2.74, respectively, for G0 through G4, and 2.27 for the offspring in G5. It is apparent that the *µ*_*g*_ parameter corresponding to the covariate **J** is the mean of the founder population and not the mean breeding value of selected individuals. In analysis JC with the covariates centered, the intercept *β* is the value of *g_i_* when (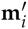 − 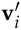) ***α*** = 0, which is the case when 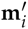 = 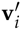. The estimate 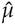 was 17.33 in this analysis, but 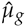 remained about the same value, namely 9.05. This shows that 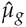 has the same interpretation whether the entire matrix of observed and imputed genotypes is centered or not. In neither case does it represent the mean breeding value of selected individuals.

#### 2. Bias

Table 3 shows the regression coefficients of TBV on (G)EBV for *h*^2^ = 0.5 and *µ*_*α*_ = 0.2, the same scenarios represented in Table 2. The regression coefficients of TBV on GEBV for each scenario were close to 1 with very low SE, which indicates the genomic predictions exhibited almost no bias. The differences in regression coefficients between analyses were very small, but the marker panel comprising only markers were biased upwards whereas the marker panels that included causal mutations were biased slightly downwards.

**Table 3.** Regression coefficients (±*SE*_*s*_) of TBV on (G)EBV^*a*^

| Genotype Data<sup>b</sup> | Analyses<sup>c</sup> |  |  |  |  |
| --- | --- | --- | --- | --- | --- |
|  | JC | J | C | N | PBLUP |
| 50 QTL + 50 Markers | 1.06 ± .01 | 1.05 ± .01 | 1.03 ± .01 | 1.03 ± .01 | - |
| 50 QTL Only | 1.05 ± .01 | 1.05 ± .01 | 1.04 ± .01 | 1.04 ± .01 | - |
| 50 Markers Only | 0.82 ± .02 | 0.82 ± .02 | 0.82 ± .02 | 0.82 ± .02 | - |
| No Genotypes | - | - | - | - | 0.92 ± .02 |

<sup>a</sup> Average Regression coefficients of true breeding value (TBV) on (genomic) estimated breeding values from 10 replications validated in Generation 5, comprising 8,000 individuals with genotypes but no phenotypes. The true QTL effects were sampled from a Normal distribution with mean $\mu_a = 0.2$ and scaled to simulate a trait with a heritability 0.5.
<sup>b</sup>The analyses were based on fitting covariates for only 50 QTL, only 50 markers, or both 50 QTL and 50 markers.
<sup>c</sup>J: includes a covariate for $\mu_g$ in the model, C: entire matrix of imputed and observed genotype covariates centered, JC: both J and C, N: neither J or C, and PBLUP: pedigree-based BLUP.

### Sensitivities to trait heritability

Accuracy of PBLUP increased with heritability, as expected (Table 4). Genomic predictions using panels that include causal mutations were near perfect when *µ*_*g*_ was included in the model. These high accuracies are a reflection of these phenotypes being influenced by only 50 QTL and there being a large training dataset. Accuracy was reduced when *µ*_*g*_ was not fitted in the model. There was no advantage in terms of accuracy to centering the covariates but MCMC mixing may have been improved although this was not investigated.

**Table 4.** Correlations (%, ± *SE*_*s*_) between TBV and (G)EBV^*a*^ for alternative analyses^*b*^ for different heritabilities

| Heritabilities | Analyses |  |  |  |  |
| --- | --- | --- | --- | --- | --- |
|  | JC | J | C | N | PBLUP |
| $h^2 = 0.1$ | 94.91 ± .00 | 94.89 ± .00 | 89.44 ± .01 | 89.46 ± .01 | 30.29 ± .01 |
| $h^2 = 0.3$ | 97.93 ± .00 | 97.97 ± .00 | 92.55 ± .01 | 92.53 ± .01 | 37.61 ± .01 |
| $h^2 = 0.5$ | 98.35 ± .00 | 98.41 ± .00 | 92.20 ± .00 | 92.18 ± .00 | 41.53 ± .00 |

<sup>a</sup> Average correlation between true breeding value (TBV) and (genomic) estimated breeding values from 10 replications validated in Generation 5, comprising 8,000 individuals with genotypes but no phenotypes. The true QTL effects were sampled from a Normal distribution with mean $\mu_a = 0.2$ and scaled to simulate a trait with a heritabilities 0.1, 0.3 or 0.5.
<sup>b</sup>J: includes a covariate for $\mu_g$ in the model, C: entire matrix of imputed and observed genotype covariates centered, JC: both J and C, N: neither J or C, and PBLUP: pedigree-based BLUP. Covariates were fitted for both 50 QTL and 50 Markers.

### Effect of mean QTL effect (µ_α_ = 0 vs µ_α_ = 0.2)

We had hypothesized that the impact of omitting *µ*_*g*_ from the model will be greatest when *µ*_*g*_ departs significantly from 0 which is more likely to occur when *µ*_*α*_ departs from 0. For that reason our base simulation used *µ*_*α*_ = 0.2. Results are shown in Table 5 for the panel including QTL and markers with *h*^2^ = 0.5 for *µ*_*α*_ = 0.2 compared to *µ*_*α*_ = 0. These results confirmed the benefit of fitting *µ*_*g*_ was greatest when *µ*_*α*_ = 0.2 but there was still an advantage to fitting *µ*_*g*_ when *µ*_*α*_ = 0. That advantage is likely to erode as the number of QTL increases. Changing the mean QTL effect had no impact on bias, except for a slight influence on PBLUP.

**Table 5.** Accuracy^*a*^ and bias^*b*^ of genomic prediction (±*SE*_*s*_) for alternative QTL distributions^*c*^ and analyses^*d*^

| Substitution<br>Effects | Analyses |  |  |  |  |
| --- | --- | --- | --- | --- | --- |
|  | JC | J | C | N | PBLUP |
| **Correlations (%)** |  |  |  |  |  |
| $\mu_\alpha = 0$ | 98.63 ± .00 | 98.66 ± .00 | 97.31 ± .00 | 97.31 ± .00 | 41.87 ± .01 |
| $\mu_\alpha = 0.2$ | 98.35 ± .00 | 98.41 ± .00 | 92.20 ± .00 | 92.18 ± .00 | 41.53 ± .00 |
| **Regression Coefficient** |  |  |  |  |  |
| $\mu_\alpha = 0$ | 1.07 ± .03 | 1.07 ± .03 | 1.06 ± .02 | 1.06 ± .02 | 0.95 ± .03 |
| $\mu_\alpha = 0.2$ | 1.06 ± .01 | 1.05 ± .01 | 1.03 ± .01 | 1.03 ± .01 | 0.92 ± .02 |

<sup>a</sup> Accuracy was quantified using the average correlation between true breeding value and (genomic) estimated breeding values from 10 replications validated in Generation 5, comprising 8,000 individuals with genotypes but no phenotypes.
<sup>b</sup> Bias was quantified using the average regression coefficients of true breeding value on (genomic) estimated breeding values from 10 replications.
<sup>c</sup> The true QTL effects were sampled from Normal distributions with mean $\mu_\alpha = 0$ or $\mu_\alpha = 0.2$ and scaled to simulate a trait with a heritability 0.5.
<sup>d</sup> J: includes a covariate for $\mu_g$ in the model, C: entire matrix of imputed and observed genotype covariates centered, JC: both J and C, N: neither J or C, and PBLUP: pedigree-based BLUP. Covariates were fitted for both 50 QTL and 50 markers.

### Effect of more QTL and markers (100 SNP vs 1,000 SNP)

We had hypothesized that the improvement of accuracy from adding an extra covariate for *µ*_*g*_ will reduce as the number of QTL increases because *µ*_*g*_ is likely to be closer to zero for a trait that is more polygenic. Table 6 shows that PBLUP was largely unaffected by changes to genetic architecture but the accuracy of genomic prediction declined slightly as the number of QTL increases. This reflects the fact that precision of estimating QTL effects is greater when the effects are large and polygenic traits with more QTL must have on average smaller effects when compared at the same genetic variance. The benefit of fitting *µ*_*g*_ in the model was virtually eliminated when the number of substitution effects to estimate increases from 100 to 1,000. In contrast to the results of accuracy, there was no impact of QTL number on bias. Centering had no impact on accuracy or bias.

**Table 6.** Accuracy^*a*^ and bias^*b*^ of genomic prediction (±*SE*_*s*_) for different numbers of QTL^*c*^ and alternative analyses^*d*^

| Numbers of QTL | Analyses |  |  |  |  |
| --- | --- | --- | --- | --- | --- |
|  | JC | J | C | N | PBLUP |
| **Correlations (%)** |  |  |  |  |  |
| **50 QTL + 50 Markers** | 98.35 ± .00 | 98.41 ± .00 | 92.20 ± .00 | 92.18 ± .00 | 41.53 ± .00 |
| **500 QTL + 500 Markers** | 95.49 ± .00 | 95.48 ± .00 | 94.87 ± .00 | 94.86 ± .00 | 41.48 ± .00 |
| **Regression Coefficient** |  |  |  |  |  |
| **50 QTL + 50 Markers** | 1.06 ± .01 | 1.05 ± .01 | 1.03 ± .01 | 1.03 ± .01 | 0.92 ± .02 |
| **500 QTL + 500 Markers** | 1.04 ± .01 | 1.04 ± .01 | 1.04 ± .01 | 1.04 ± .01 | 0.92 ± .01 |

<sup>a</sup> Accuracy was quantified using the average correlation between true breeding value and (genomic) estimated breeding values from 10 replications validated in Generation 5, comprising 8,000 individuals with genotypes but no phenotypes.
<sup>b</sup> Bias was quantified using the average regression coefficients of true breeding value on (genomic) estimated breeding values from 10 replications.
<sup>c</sup> The true effects for 50 or 500 QTL were sampled from a Normal distribution with mean $\mu_\alpha = 0.2$ and scaled to simulate a trait with a heritability 0.5.
<sup>d</sup> J: includes a covariate for $\mu_g$ in the model, C: entire matrix of imputed and observed genotype covariates centered, JC: both J and C, N: neither J or C, and PBLUP: pedigree-based BLUP. Covariates were fitted for either 50 QTL and 50 markers or 500 QTL and 500 markers.

### Centering using the entire matrix of genotype covariates or only the observed genotype covariates

Table 7 shows the accuracies and regression coefficients of TBV on G(EBV) for the genotype panel with 50 markers, *h*^2^ = 0.5 and *µ*_*α*_ = 0. The analyses were performed after centering: the entire matrix of imputed and observed genotype covariates (**M**\* = **M** −**11**′**M**); only observed genotype covariates (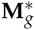 = **M**_g_ − **11**′**M**_*g*_), which is the type of centering done in single-step genomic best linear unbiased prediction (GBLUP); or not centering the covariates (**M**\* = **M**). The accuracy of any genomic analyses were about 8% higher than that one based on covariates centered as 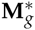 but without **J** (model C*). However, when **J** was included in the model with covariates centered as 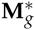, the accuracy of prediction was markedly improved.

**Table 7.** Accuracy^*a*^ and bias^*b*^ of genomic prediction (±*SE*_*s*_) when centering for all genotypes or observed genotypes^*c*^

| Analyses | Correlations (%) | Regression Coefficient |
| --- | --- | --- |
| JC | 73.36 ± 0.03 | 0.89 ± 0.02 |
| J | 73.37 ± 0.03 | 0.89 ± 0.02 |
| C | 73.07 ± 0.03 | 0.89 ± 0.02 |
| N | 73.07 ± 0.03 | 0.89 ± 0.02 |
| JC* | 73.38 ± 0.03 | 0.89 ± 0.02 |
| C* | 65.30 ± 0.03 | 1.32 ± 0.06 |
| PBLUP | 41.87 ± 0.01 | 0.95 ± 0.03 |

<sup>a</sup>Accuracy was quantified using the average correlation between true breeding value and (genomic) estimated breeding values from 10 replications validated in Generation 5, comprising 8,000 individuals with genotypes but no phenotypes. The true QTL effects were sampled from Normal distributions with mean $\mu_\alpha = 0$ and scaled to simulate a trait with a heritability 0.5.
<sup>b</sup>Bias was quantified using the average regression coefficients of true breeding value on (genomic) estimated breeding values from 10 replications.
<sup>c</sup>**J**: includes a covariate for $\mu_g$ in the model, **C**: entire matrix of imputed and observed genotype covariates centered, **JC**: both **J** and **C**, **N**: neither **J** or **C**, **C\***: only observed genotype covariates centered, **JC\***: both **J** and **C\***, and **PBLUP**: pedigree-based BLUP. Covariates were fitted for 50 markers.

As explained previously, *µ*_*g*_ = **k**′***α***, where **k**′ is the expected value of the covariates in the founders, will tend to zero for the marker panel that does not include QTL even with *µ*_*α*_ ≠ = 0. However, even if *µ*_*α*_ = 0, in a population undergoing selection when selected individuals are genotyped, 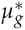 = *µ*_*g*_ − **v**′ ***α*** ≠ = 0, where **v**′ is the expected value of the observed genotype covariates. In this study, selection was used to increase the mean of the trait. Thus, 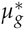 is expected to be negative because most of the genotyped individuals were from G5, whereas *µ*_*g*_ is expected to be zero. The negative estimate of 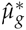 from 10 replicates of the JC* analysis, −2.84, confirms that 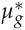 < 0. On the other hand, the mean of 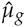 from 10 replicates of the JC analysis was −0.23. This explains why fitting **J** in the model improved the accuracy of genomic prediction when covariates were centered as in single-step GBLUP.

Fernando et al. (Fernando *et al*. 2014) found that centering using 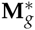 improves the accuracy without **J** in the model when the population was not under selection and the genotyped individuals were unselected. In that study, mating was random with no selection, so the allele frequency means of the genotyped individuals were a reasonable approximation of the allele frequency means in the founder population. In contrast, our simulation here shows that centering using 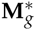 can reduce the accuracy when the population is under selection, unless **J** is fitted in the model.

In single-step GBLUP, the observed genotypes are commonly centered by subtracting their mean and used to construct a genomic relationship matrix, such as using the first method proposed by VanRaden (VanRaden 2008). Using that genomic relationship matrix in the single-step GBLUP formula in Aguilar et al. (Aguilar *et al*. 2010) does not account for **J**. This was recognised in Vitezica et al (Vitezica *et al*. 2011) who proposed a modification for populations under selection that involved adding a constant to all elements of the genomic relationship matrix that they derived by equating the sum of the elements of the genomic relationship matrix to the the sum of the elements of the numerator relationship matrix. In the appendix of that paper they showed this modification is equivalent to fitting a covariate **Q** = − **J** and treating −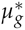 as a random effect. In addition to this modification, Christensen et al. (Christensen *et al*. 2012) proposed a multiplicative scaling to the genomic relationship matrix such that its diagonals have the same mean as the diagonals of the numerator relationship matrix. Vitezica et al. (Vitezica *et al*. 2011) claimed that −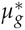 represents the mean breeding value of selected individuals, and we have confirmed here that this is true provided the observed genotype covariates are centered by their mean.

Most populations are under natural or artificial selection. In many cases, genotypes are only available on selected individuals. In single-step genomic analysis that combine genotyped and non-genotyped individuals in a joint analysis, the mean of observed genotypes are available for centering. If the observed genotypes include QTL, the accuracy of genomic prediction can be severely compromised, unless the **J** covariate is fitted in the model. If the observed genotypes are only markers, the accuracy of genomic prediction may not necessarily be improved by fitting **J** in the model, but it doesn’t do any harm. However, if centering is applied only to the observed genotypes, which is the type of centering used in single-step GBLUP, accuracy could be severely compromised, unless the **J** covariate is fitted in the model or an equivalent approach is adopted.

## Acknowledgements

The authors are grateful to Bruce L. Golden and Hao Cheng for assistance in the implementation of SSBR models, and to Jack C. M. Dekkers for his constructive comments in design of simulation. This work was supported by the US Department of Agriculture, Agriculture and Food Research Initiative National Institute of Food and Agriculture Competitive grant no. 2015-67015-22947.

## Appendix

Here we show that inference about ***α*** does not depend on how the genotypes are coded. The marker effects model can be described by the following general model:

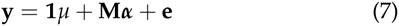

where **y** is a vector of observed phenotypes, **1** is a vector of 1s, *µ* is a general mean, **M** is a matrix of marker covariates, coded 0, 1, 2, which represents the number of copies of the A allele, ***α*** is a vector of random substitution effects of markers, and e is a vector of residuals. Henderson’s mixed model equations (MME) that correspond to equation (7) are:

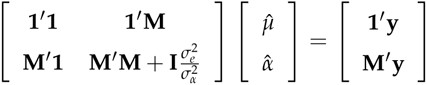

where 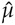 is the best linear unbiased estimate of the mean, and 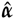 is the best linear unbiased predictor of the vector of random substitution effects of all markers. Now we can eliminate 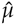 from the equations for 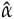, by subtracting from those equations the equation for 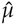 pre-multiplied by 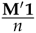. Then, the MME are transformed to:

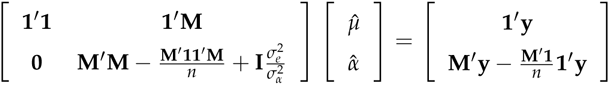

and substituting **1**′ **1** = *n* and **1**′ **M** = *n*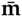′ and its transpose, the transformed MME become:

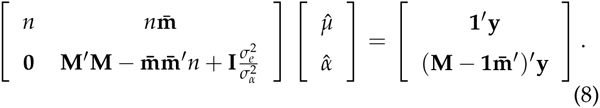

where 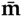′ is the row vector of column means of **M** as in 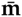′ = 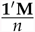.

Now, consider the coding obtained by centering the marker genotypes as **M** − **1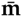**′. Then the model can be written as:

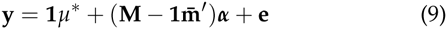

where *µ*\* = *µ* + 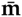′***α***. The MME that correspond to equation (9) are:

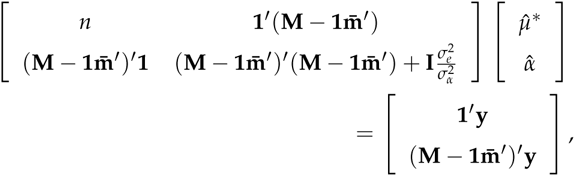

but **1**′(**M** − **1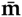**′) = *n*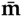*′ − n*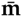′ = **0**′, and, similarly, its transpose is (**M** − **1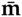**′)′ **1** = **0**. Then the MME become:

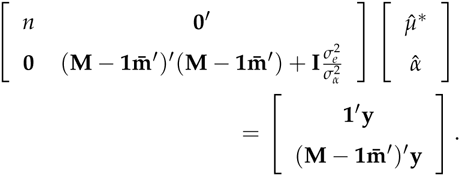

Expanding (**M** − **1** 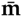′)′(**M** − **1** 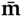′) gives **M**′ **M** −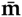**1**′ **M** − **M**′ **1** 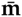′ + 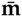**1**′ **1** 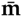′, but because **1**′ **M** = *n*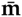′, **M**′ **1** = *n*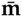, and **1**′ **1** = *n*, as previously shown, the second term, 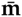**1**′ **M** = 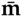 *n*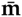′, which is equal to the last term in the expansion. Thus, the MME become:

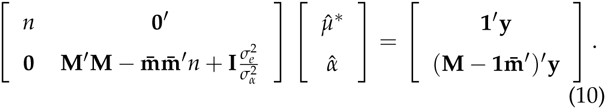

The equations for 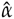 in (8) and (10) are identical, and this proves that centering with **m**′ doesn’t affect inference about ***α***.

Now suppose an arbitrary vector **v**′ is used to transform the genotypes as (**M** − **1v**′). The the model becomes:

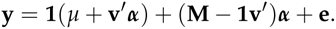

Adding and subtracting **1** 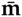′***α***, the above equation can be written as:

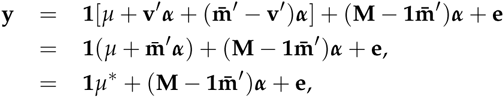

which is identical to equation (9), proving inference about ***α*** does not depend on how the genotypes are coded.

